# Fiber photometry analysis of spontaneous dopamine signals: The z-scored data are not the data

**DOI:** 10.1101/2025.02.19.639080

**Authors:** Conner W Wallace, Clare Y Slinkard, Rachael Shaughnessy, Katherine M Holleran, Samuel W Centanni, Christopher C Lapish, Sara R Jones

**Author notes:** Corresponding author: Sara R Jones.

## Abstract

Fluorescent sensors have revolutionized the measurement of molecules in the brain, and the dLight dopamine sensor has been used extensively to examine reward- and cue-evoked dopamine release, but only recently has the field turned its attention to spontaneous release events. Analysis of spontaneous events typically requires evaluation of hundreds of events over minutes to hours, and the most common method of analysis, z-scoring, was not designed for this purpose. Here, we compare the accuracy and reliability of three different analysis methods to identify pharmacologically induced changes in dopamine release and uptake in freely moving C57BL/6J mice. The D1-like receptor antagonist SCH23390 was used to prevent dLight sensors from interacting with dopamine in the extracellular space, while cocaine was used to inhibit uptake and raclopride to increase release of dopamine in the nucleus accumbens. We examined peak-to-peak frequency, peak amplitude, and width, the time spent above an established cutoff. The three methods were 1) the widely-used “Z-Score Method”, which automatically smooths baseline drift and normalizes recordings using signal-to-noise ratios, 2) a “Manual Method”, in which local baselines were adjusted manually and individual cutoffs were determined for each subject, and 3) the “Prominence Method” that combines z-scoring with prominence assessment to tag individual peaks, then returns to the preprocessed data for kinetic analysis. First, SCH23390 drastically reduced the number of signals detected as expected, but only when the Manual Method was used. Z-scoring failed to identify any changes, due to its amplification of noise when signals were diminished. Cocaine increased signal width as expected using the Manual and Prominence Methods, but not the Z-Score Method. Finally, raclopride- induced increases in amplitude were correctly identified by the Manual and Prominence Methods. The Z-Score Method failed to identify any of the changes in dopamine release and uptake kinetics. Thus, analysis of spontaneous dopamine signals requires assessment of the %ΔF/F values, ideally using the Manual Method, and the use of z-scoring is not appropriate.

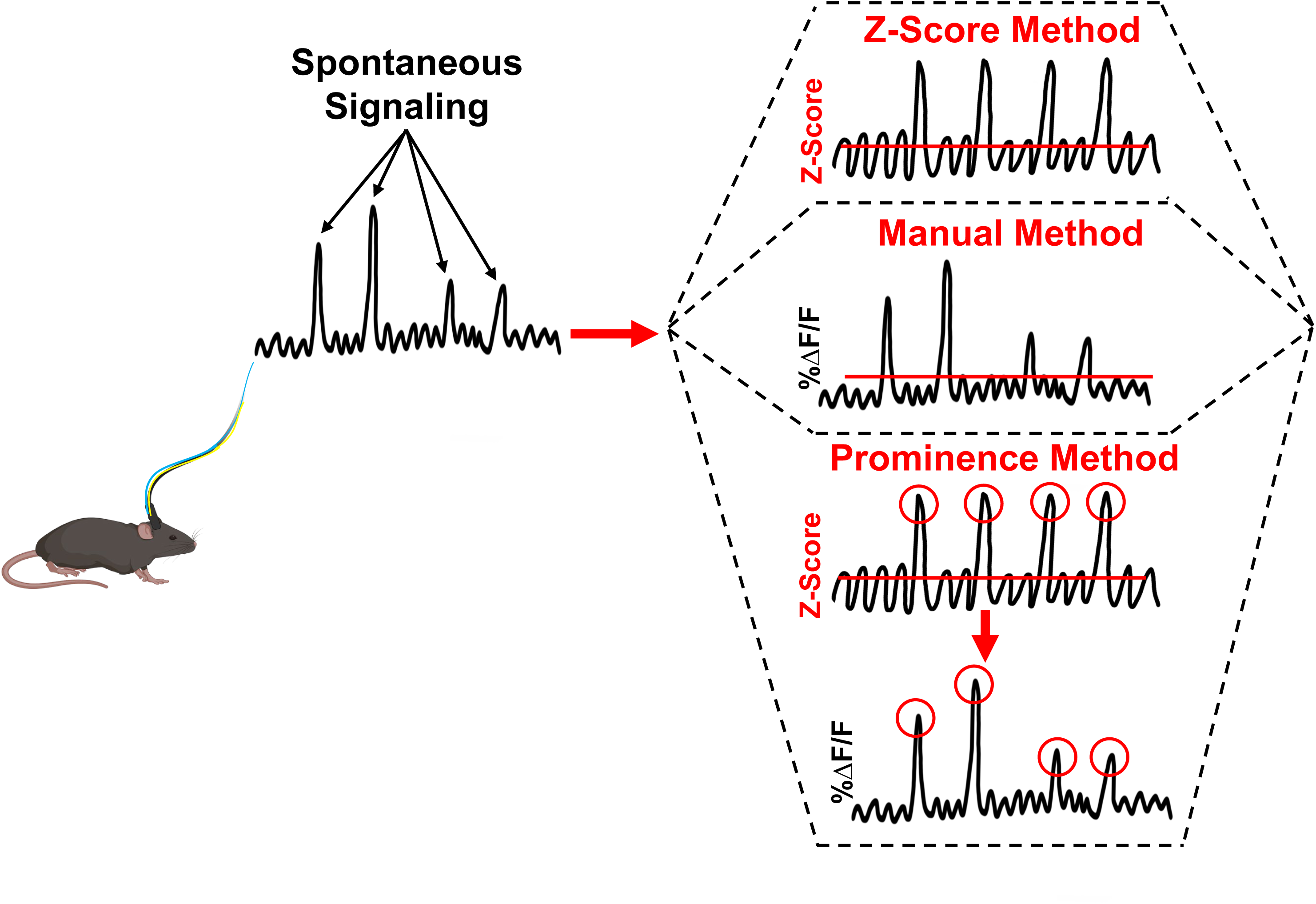

## 1. Introduction

The measurement of real-time neuronal activity in freely moving animals was revolutionized over 20 years ago by the use of fiber photometry (FP) paired with GCaMP, a highly sensitive fluorescence-based calcium sensor with a time resolution of milliseconds [1–4]. Since then, sensors have been developed for many different molecules, including multiple neurotransmitters and neuromodulators, and the list is growing rapidly (for an excellent review, see [5]). Several recent papers on FP [5–8] have sought to create a standardized guide for FP acquisition and analysis. While open-source options for FP analysis have been published [9, 10], there is no true standard in the field other than the use of z-scoring to analyze behavior-linked events, whether by open-source methods or those developed in-house. However, the vast majority of dopamine (DA) signals observed in our laboratory and others [11, 12] are not directly linked to behaviors or external stimuli, but appear to arise spontaneously. With the extensive use of FP to measure neural activity in real time, there remains a lack of standardized analysis techniques to assess spontaneous signals and their kinetics.

Importantly, we found the use of z-scoring to normalize the baseline and then derive FP signals, the “gold standard” for analysis of discrete event-related FP signals, was completely inappropriate for analysis of spontaneous signals.

This paper will focus on how to analyze spontaneously arising, endogenous DA release events measured using the dLight DA sensor [13] coupled with fiber photometry. The popularity of DA sensors like dLight and GRAB_DA_ [6, 8, 14] has exploded, with widespread adoption of these sensors for *in vivo* DA measurements. The vast majority of DA-focused FP experiments have focused on fluctuations around discrete events such as the presentation of cues [15, 16], rewards or noxious stimuli [17, 18], or behaviors such as lever presses [19, 20], nose pokes [21, 22], reward seeking or consumption [22, 23], or approach/avoidance behavior [23–26]. For analysis of such events, it is common to use normalization through z-scoring, which determines the relative strength of the change in signal over time compared to the variability present at baseline [5]. This approach is well validated for behavioral events and is necessary due to photobleaching, because the signal will drift throughout a prolonged recording, requiring a new baseline to be established close to each behavioral event of interest. This is followed by analysis of a narrow timeframe around that event.

There are several caveats to analyzing *spontaneous* signals recorded using FP. One issue was at least partly addressed in a recent publication: normalization of entire recordings through z-scoring can “remove variation of experimental interest” [5]. The authors specifically called for a direct and rigorous comparison of various analysis methods for different applications. The implication here is that experimental effects on spontaneous signals could be diminished through z-scoring if the comparison or control recordings were taken at a distant point in time, because there is no reference for the current recording by which to assess *changes* in variability (on which the z-score value is dependent). Indeed, we have discovered that z-scoring does not work for assessing spontaneous signaling in longitudinal studies when signal kinetics will change session-to-session (e.g., due to drug pretreatment or behavioral interventions). Thus, we sought to determine best practices for spontaneous signal analysis by comparing the use of three distinct methods applied to the same dLight dataset.

A guiding principle to ascertain the accuracy of our analyses was the fact that the maximal achieved “peak height” of DA release is highly correlated with the amount of time required for clearance from the synapse, because more DA takes longer to be cleared by the dopamine transporter (DAT) [27–29]. To assess how underlying changes in the kinetics of real-time, spontaneous DA fluctuations can be measured by dLight, we specified a cutoff point above which signals were analyzed for amplitude (peak height), width (time for signal to rise above then fall below the cutoff), and frequency. We further derive a variable termed ‘slope’ by correlating amplitude with width. The three methods we used to analyze these kinetics were 1) smoothing and normalization through the use of z-scoring, using a standard z-score cutoff across all trials and animals (Z-score Method - ZM), 2) manual baseline adjustment of non-normalized %ΔF/F data followed by prescribing a %ΔF/F cutoff for each animal (Manual Method - ManM) and 3) a method that uses traditional z-scoring for signal detection with a standard cutoff followed by analysis of the non-normalized %ΔF/F data at the same timepoints, with signal inclusion determined by signal “prominence” compared to background noise (Prominence Method - ProM). We aimed to assess how well these distinct analysis methods performed at identifying drug effects on spontaneous DA signals.

It is common in FP studies to normalize recordings around behavioral events to obtain z-score values, which are used to quantify and compare different aspects of signaling. Therefore, we hypothesized that the ZM would perform better than the ManM at detecting drug-induced changes to the kinetics of spontaneous signals, and further that the ProM would perform the best at detecting changes in signal amplitude because it assesses peak prominence. To directly test this, we injected dLight into the nucleus accumbens (NAc) core of male and female mice. Animals were then exposed to pharmacological challenges (SCH23390 (a DA D1-like receptor (D1R) antagonist), cocaine (a DA transporter (DAT) inhibitor), raclopride (a DA D2-like receptor (D2R) antagonist), and vehicle), and dLight signals were recorded. We hypothesized that the primary effects of each drug would be a reduction in signal frequency by SCH23390, an increase in width by cocaine, and an increase in amplitude by raclopride. It was further expected that all three methods would identify these changes. In direct contrast, the ZM was unable to detect any of these primary drug effects. Analysis of the non-normalized %ΔF/F values, as with the ManM and ProM, was superior in detecting drug-induced changes in spontaneous signaling. Below, we describe the effort to accurately assess alterations to the kinetics of real-time DA signaling using dLight. Ultimately, we posit that z-scoring cannot be used for the final analysis of spontaneous DA signals.

## 2. Results and Discussion

### 2.1. Three methods were developed around a central premise of adjusting the baseline and applying a cutoff to assess variables

Figure 1 is a brief introduction to the three methods. Preprocessing included the generation of either z-scored or non-normalized %ΔF/F traces, followed by cutoff selection and final analysis. In the Z-Score Method (ZM), the isosbestic control signal (excited at 405 nm) and experimental channel (excited at 465 nm) were smoothed and fitted using linear regression, normalized into z-score values, then subtracted to generate a z-scored ΔF/F trace (Figure 1A). A standardized approach [16] was used to determine a z-score cutoff of z = 2.6, which was applied across all animals and trials, and z-scored values were analyzed. In the Manual Method (ManM), the raw isosbestic channel was subtracted from the experimental channel. Next, %ΔF/F traces were input into pCLAMP Clampfit software without any automatic processes like regression, smoothing, or normalization through z-scoring. After manually adjusting the baseline for signal drift, a %ΔF/F cutoff was defined for each animal during drug-naïve, saline control trials, and this cutoff was maintained within animals across drug trials (Figure 1B). In the third and final method (Prominence Method (ProM); Figure 1C), the experimental channel was adjusted for bleaching and artifacts by normalizing it into z-scores and fitting a 100 s moving-average filter. A standard z-score cutoff across all animals and trials was applied to *detect* signals. The detected peaks were timestamped, after which they were assessed for prominence within the non-normalized %ΔF/F trace. Only signals that were sufficiently large compared to the signaling immediately around each detected peak were included. Finally, the same timestamps generated during z-scoring were used to return to the non-normalized %ΔF/F trace to pull values for analysis.

**Figure 1.**
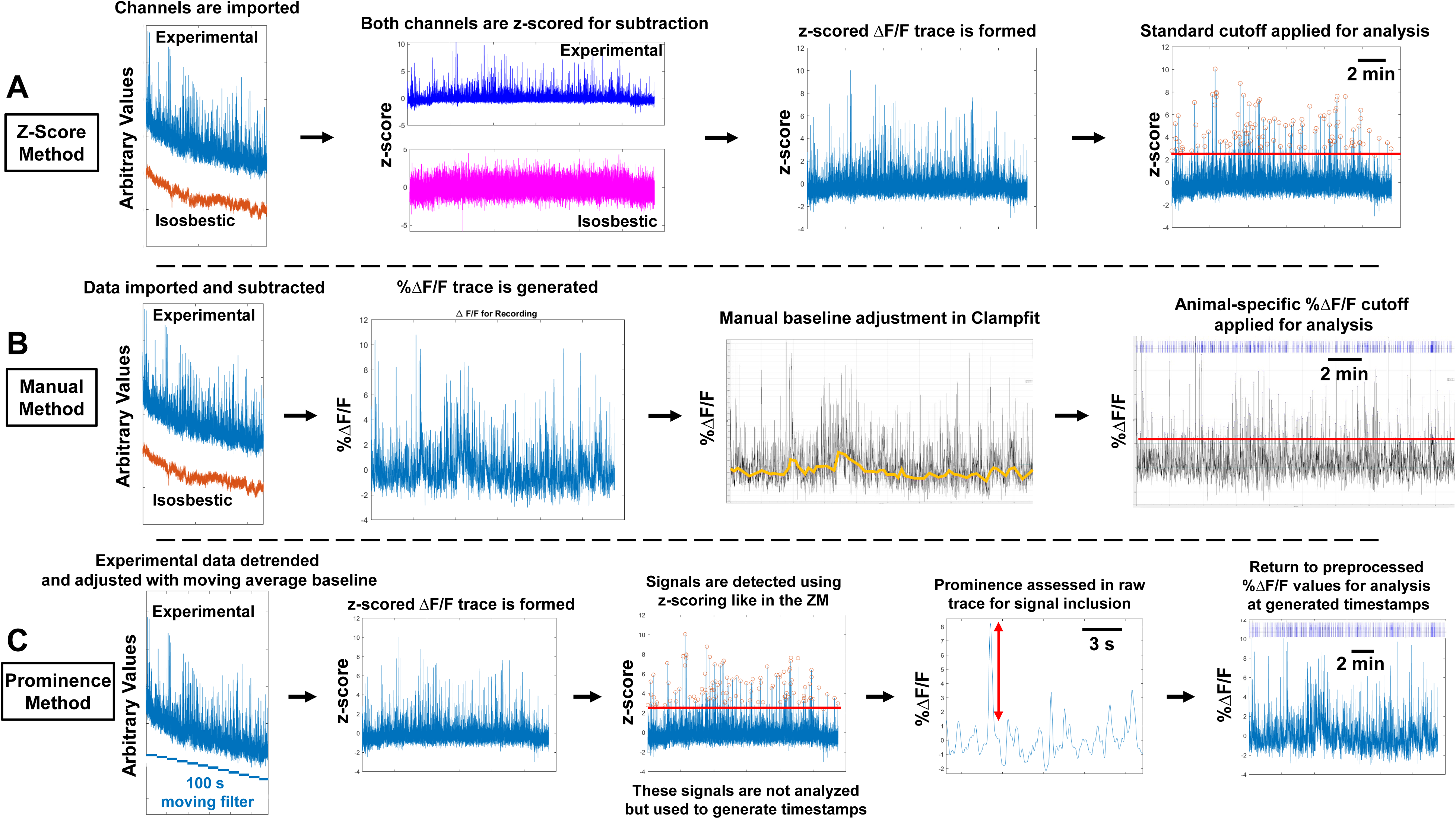
The three analysis methods are introduced, which are drastically different in efficacy of detecting pharmacological effects. The central premise of analysis is applying a cutoff to the recording, above which variables are assessed, including amplitude, width, and frequency. (A) The Z-Score Method (ZM) smooths and normalizes traces and applies a standard z-score cutoff. (B) The Manual Method (ManM) inputs the non-normalized, preprocessed %ΔF/F trace into Clampfit for manual baseline adjustment, and fits a specific cutoff to each animal, which is maintained over pharmacological conditions. (C) Finally, the Prominence Method (ProM) uses z-scoring for signal detection, followed by assessing the prominence of each signal compared to its immediate surroundings to decide which signals to include, then uses the corresponding data in the %ΔF/F trace for analysis.

It is important to note that both the ManM and ProM used preprocessed %ΔF/F values for *analysis,* whereas the ZM used z-scored values. Further, the absolute minimum amplitude values detected by the ManM and ZM were determined by the definitive cutoff applied with each method, whether using a %ΔF/F or z-score cutoff, respectively. For example, with a ΔF/F cutoff of 2.0% using the ManM, no values below 2% can be included. In contrast, when using the ProM, the minimum amplitude values may be different between sessions by using prominence to decide which signals to include, because the variability inherent to each session might differ over time and between pharmacological conditions.

### 2.2. Blocking the dLight sensor with a D1 receptor antagonist abolished detection of spontaneous signals, an effect shown only through use of the ManM

A DA D1-like receptor (D1R)-antagonist was used to ensure spontaneous dLight signals were a result of activating the dLight receptors, rather than spurious noise.

SCH23390 blocks D1Rs, including the D1Rs conjugated to GFP in dLight, which allows the sensor to fluoresce in response to DA. This means that SCH23390 should almost completely reduce detection of spontaneous signals above a given cutoff, reducing the signal frequency, amplitude, and width. Results are presented for the primary hypothesized effect, a reduction in frequency. Given that there is not an established method of assessing spontaneous dLight signals, we initially turned to z-score normalization, as is standard for assessing behavior-associated release events.

However, a reduction in dLight signal frequency in response to SCH23390 was not observed using the ZM (t_5_ = 0.6447; p=0.55; Figure 2A), causing us to re-evaluate our choice of analysis methods for spontaneous signals. Using the ManM, a simple two-tailed t-test showed that SCH23390 reduced signal frequency (t_5_ = 4.831; saline: 0.28 ± 0.046 signals/s; SCH23390: 0.022 ± 0.015 signals/s; p<0.01; Figure 2B). The ProM, which uses z-scoring to initially identify signals like the ZM, also did not identify a difference in frequency (t_5_ = 2.322; p=0.0679; Figure 2C). To confirm this difference between methods, we next calculated the average frequency of saline trials and set it to a baseline of 0% to determine the percent change due to SCH23390. One-way analysis of variance (ANOVA) identified a significant effect of analysis method (F_(1.117,5.586)_ = 60.04; p<0.001). Post-hoc analysis showed the ManM resulted in a stronger reduction in signal frequency (-92 ± 5%) than the ZM (-5 ± 3%; p<0.0001) and the ProM (+28 ± 14%; p<0.001; Figure 2D). This suggests that noise is amplified using the ZM and ProM under a condition when the dLight receptor is blocked because the entirety of these longitudinal recordings, which were taken on separate days, were smoothed and normalized using these methods. Such amplification of noise could lead to erroneous detection of signal. This finding also highlights the possibility that the adjustments inherent to these analysis methods may falsely obscure or diminish the observation of real effects of interventions that influence spontaneous events over the course of the recording period, such as a drug pretreatment.

**Figure 2.**
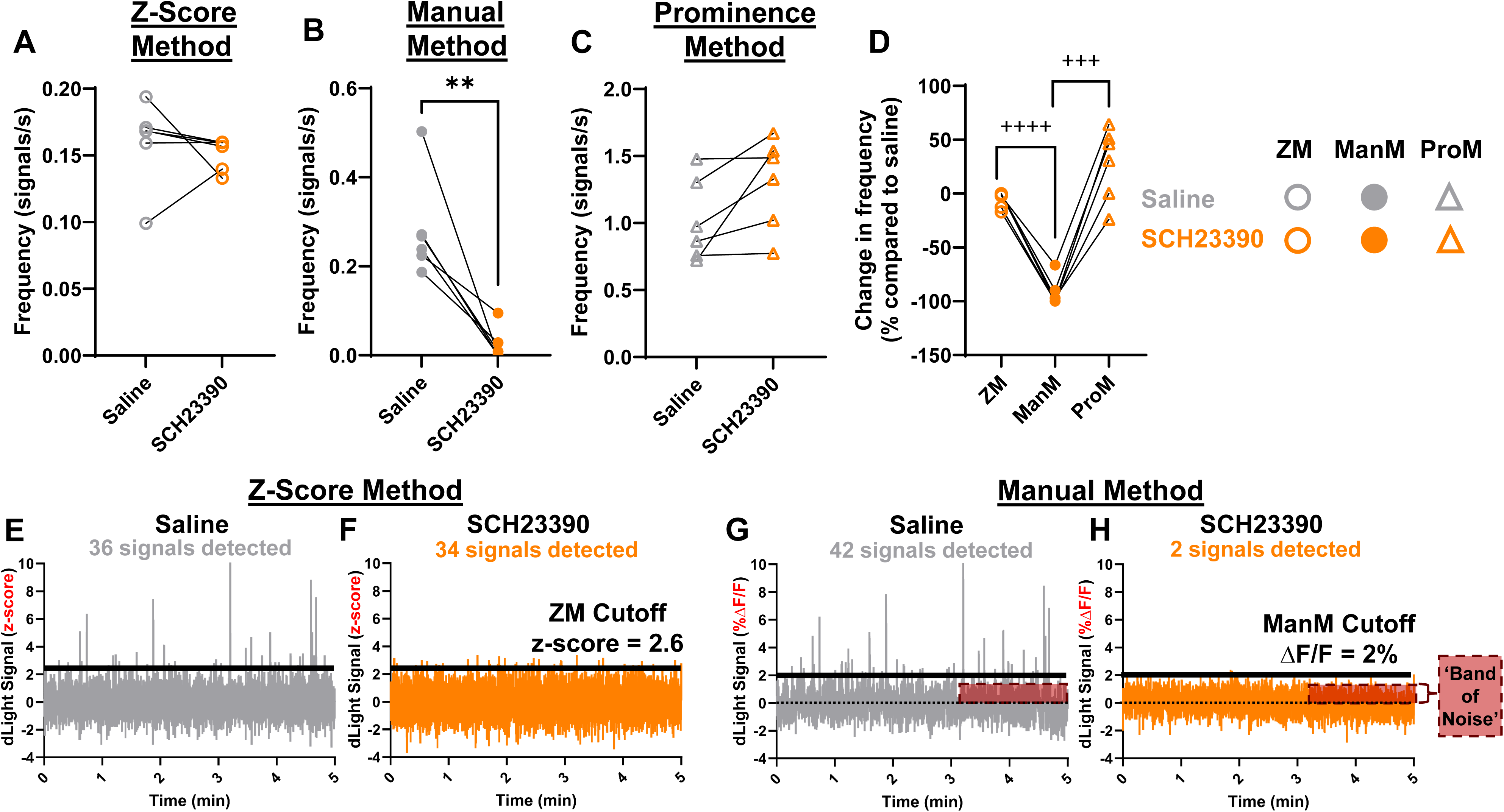
Normalization through z-scoring prevents detection of a reduction in frequency of spontaneous signals induced by blockade of the dLight receptor. SCH23390, a D1R antagonist, blocks activation of the D1R-based dLight receptor, which should fully attenuate the frequency of detected signals compared to saline trials. (A) This difference was not detected using the Z-Score Method (ZM) between saline ( ○ ) and SCH23390 ( ○ ) trials. (B) Using the Manual Method (ManM), SCH23390 ( ● ) reduced frequency of detected signals compared to saline ( ● ). (C) However, this change was not detected using the Prominence Method (ProM) between saline ( Δ ) and SCH23390 ( Δ ) trials. (D) Results from each method are directly compared, showing that the ManM performed better than the ZM and ProM at detecting a reduction in frequency by SCH23390. (E-F) Visual examples are shown highlighting the issues with the ZM. Five-minute traces are shown as analyzed by the ZM with sessions from the same animal during (E) saline and (F) SCH23390, highlighting that a significant number of signals were identified during the SCH23390 trial. Artificial noise was amplified in the absence of signals that result from DA activation of the dLight receptor. (G-H) The same traces are shown as analyzed by the ManM. Only 2 signals were detected. (**, p<0.01; difference between saline and drug) (+++, p<0.001; ++++, p<0.0001; difference between methods).

To present a visual example of this issue, we present 5-minute traces of the dLight recordings for the saline control and SCH23390 sessions within the same animal in Figure 2. When analyzed using the ZM, several spontaneous signals were detected during the saline control session (Figure 2E), as expected. However, using a z-score cutoff of z = 2.6, which was used to generate values in Figure 1 and was applied across animals, many signals were also detected during the SCH23390 session (Figure 2F).

This is a restrictive cutoff, because a z-score/p-value table shows high confidence at z = 2.6 (p<0.01) that a difference from a control condition, or the baseline, was observed. In contrast, traces from the same saline and SCH23390 sessions from the same animal as above (Figure 2G-H, respectively), which have only been preprocessed into %ΔF/F values according to the ManM, show that most (95%) of the spontaneous dLight signals were blocked by the D1R antagonist. Thus, z-scoring blunts detection of pharmacologically induced, within-animal differences in signal frequency when analyzing two separate dLight recordings, whereas the ManM appropriately detected a reduction in frequency due to SCH23390.

We showed empirically that normalizing entire dLight sessions using a standard z-score cutoff impaired the ability to detect the reduction in signal frequency induced by SCH23390 compared to the ManM that assessed non-normalized data using a %ΔF/F cutoff. This is particularly shocking because the original purpose of the D1R antagonist experiment was to block changes in fluorescent signals by preventing DA activation of dLight in order to increase our confidence that the spontaneous signals we observed were valid, and not just spurious noise events. Thus, the ZM cannot be used to analyze signal frequency under a condition that broadly diminishes the presence of typically abundant spontaneous signals, because it increases the possibility of Type I error, or detecting false positive results.

Normalization and smoothing are useful tools for conducting statistical tests and in presenting data [30, 31]. The cutoff for the ZM, z = 2.6, reflects a high-confidence p-value (p<0.01), meaning each signal can be considered an “outlier” from the baseline, or, in other words, a ‘real’ DA signal resulting from activation of the dLight receptor [13]. However, most common normalization procedures for statistical testing operate under the assumption that two samples for comparison are representative of their population and share similar variability [32–34]. Based on the lack of effect of the ZM, we propose that real spontaneous signals, rather than noise, contribute to a significant degree to what is considered the “baseline” (what occurs under the cutoff), which will affect the z-scoring required to conduct the ZM and analyze signals. Pharmacological effects and changes over time can influence baseline patterns of fluorescence. Therefore, it appears that z-scoring does indeed diminish variability of experimental interest, as previously proposed [5], by erroneously normalizing signals based on baseline variability leading to diminished ability to assess effects on the bulk of signals detected above the cutoff.

### 2.3. Principles underlying assessment of DA release and uptake *in vivo* are confirmed in an *ex vivo* dataset

As stated above, one of the guiding principles of our kinetic analyses was that the peak height of DA release is related to the time required for uptake [27–29]. We wanted to reinforce that fact using *ex vivo* fast-scan cyclic voltammetry (FSCV) to assess stimulated DA release, a highly validated [27, 35–40] technique from our lab, using similar kinetic parameters described for our dLight data (specifically, peak height (μM DA) for amplitude and T_80_ (s) for width). During baseline recordings in the NAc core, using coronal brain slices taken from C57BL/6J mice, the peak height of DA release correlated with T_80_, the time in seconds required for 80% decay of the signal from peak height (r = 0.4172, p = 0.0244; **Figure S1A**). This supports that DA release and the time required for uptake are related. Further, use of FSCV showed that cocaine (30 µM) reduced DA release (t_12_ = 4.294; baseline: 2.045 ± 0.340 µM; cocaine: 1.035 ± 0.150 µM; p<0.01; **Figure S1B**) and increased T_80_ (t_12_ = 7.450; baseline: 1.048 ± 0.126 s; cocaine: 8.88 ± 1.076 s; p<0.0001; **Figure S1C**). We converted release and T_80_ to a slope metric, and cocaine reduced slope (t_12_ = 4.638; baseline: 2.194 ± 0.463 µM/s; cocaine: 0.143 ± 0.0291 µM/s; p<0.001; **Figure S1D**). This supports the calculation of slope as a measure of DA uptake rate using peak height and time required for clearance, and that expected drug-induced changes to this metric can be detected. The presence of this relationship in a FP dataset would validate the assessment of spontaneous DA signal kinetics using dLight. Given that we found the three methods differed in their ability to detect the predicted effect of SCH23390, we aimed to explore the ability of the three methods to detect changes in amplitude, width, and slope in the presence of systemically administered cocaine and raclopride.

### 2.4. The ZM blunted the detection of drug-induced differences in signal amplitude, width, and slope due to cocaine and raclopride

We ran experiments using systemic injection of cocaine (10 mg/kg) or raclopride (0.3 mg/kg), drugs which have very well-characterized effects on NAc DA [41–45], to assess changes in DA kinetics. Cocaine inhibits the DAT, leading to an increase in extracellular DA and greater subsequent activation of D2R autoreceptors that presynaptically inhibit release [41–43]. Thus, we expected an increase in signal width and a reduction in amplitude. In contrast, raclopride blocks presynaptic D2Rs, which increases DA release by disengaging negative feedback [43–45], leading us to hypothesize an increase in signal amplitude. To analyze these experiments, we first turned to the ZM. The ZM detected the reduction in amplitude due to cocaine (t_1194_ = 5.015; saline: 0.0 ± 1.3%; cocaine: -7.5 ± 0.76%; p<0.00001; Figure 3A), as hypothesized. However, the primary hypothesized effect of cocaine was an increase in signal width, and the ZM did not detect a change in width after Bonferroni correction (see **methods**; t_1194_ = 2.382; p=0.0174; Figure 3B). A simple linear regression analysis was run for each animal to determine slope, a measure of DA uptake using dLight [46], which we also validated using FSCV. Using the ZM, there was no difference in slope (t_6_ = 1.502; p=0.1837; Figure 3C). Thus, normalization and smoothing of the recording for both detection and quantification severely attenuated the ability to detect the primary pharmacological effect of cocaine in the NAc, to slow DA uptake. This result invalidates the use of the ZM to assess changes in DA uptake during spontaneous signaling.

**Figure 3.**
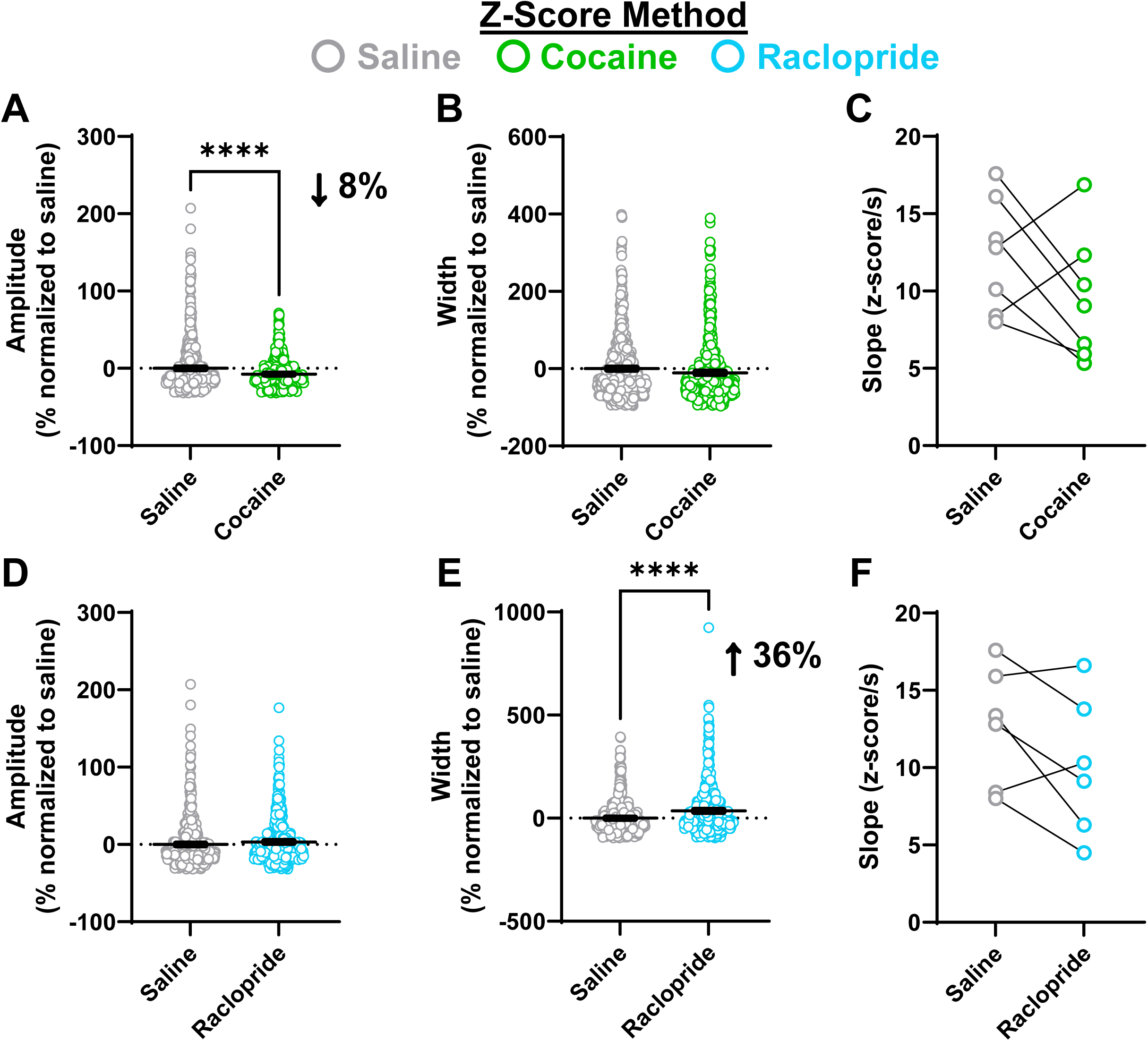
The Z-Score Method (ZM) blunts the ability to detect drug-induced changes in DA kinetics. (A-C) Analysis using the z-score normalization method (ZM) with a cutoff of z = 2.6 showed cocaine ( ○ ) reduced (A) signal amplitude but had no effect on (B) signal width or (C) slope. Data are presented as a % normalized to the average of each animal’s respective vehicle control session. The findings contrast with both other methods presented, which show an increase in signal width during cocaine sessions. (D-F) Using ZM, raclopride ( ○ ) did not affect (D) signal amplitude, but it increased (E) width without affecting (F) slope. The primary effect of raclopride herein, an increase in DA release, is obscured using the ZM. (****, p<0.00001 (Bonferroni corrected); difference between saline and drug).

When analyzed using the ZM, there was no detected change in signal amplitude due to raclopride (t_1103_ = 1.885; p=0.05972; Figure 3D), but there was an increase in signal width (t_1048_ = 7.839; saline: 0.0 ± 3.3%; raclopride: +35.9 ± 4.7%; p<0.00001; Figure 3E). Regression showed there was no difference in the slope of the correlation (t_5_ = 1.912; p=0.1141; Figure 3F). Therefore, the ZM failed to detect the primary effect of raclopride, an increase in release amplitude, and the detected effect on signal width was attenuated compared to the ManM (see **Section 2.5** for ManM results and **Section 2.7** for direct comparison). Overall, the primary expected effects of both drugs (cocaine – reduced uptake; raclopride – increased release) were significantly masked or not detected with use of a smoothing and normalization algorithm for both detection and analysis.

We measured spontaneous DA signals while animals were in the same environment with different pharmacological challenges over time. In addition to the inability to detect effects of SCH23390 to reduce dLight signal frequency (Type I error), the ZM significantly attenuated detection of reduced uptake rate by cocaine and increased DA release by raclopride. This suggests that the ZM can also promote occurrence of Type II error (false negative), because both subsequent methods discussed below detected highly significant increases in width by cocaine and release amplitude by raclopride. Therefore, a given z-score of peak release does not directly relate to %ΔF/F of dLight fluorescence *between sessions* because the ZM did not show the expected effect of raclopride to increase release compared to separate saline trials. We propose that this error not only happens for peak release, but also at the level of %ΔF/F where a signal crosses the cutoff, which would affect the assessment of width when using z-scoring and cause a lack of effect seen due to cocaine. These ideas are developed further in **Sections 2.6 and 2.7** by discussing differences between the ZM and the ProM, the latter which uses z-scoring for signal detection and %ΔF/F values for analysis. Given that endogenous DA concentrations are related to arousal, attention, affect, and motivational states [19, 47–50], and that DA can be altered by environmental [51–54], pharmacological [46, 55–57], and pathological processes [58–61], the variability in tonic DA signaling, or the ‘baseline’, will shift over time and between conditions. This suggests that using z-scoring to longitudinally measure dLight signals does not accurately detect spontaneous DA events.

### 2.5. Cocaine slowed DA uptake and raclopride increased DA release – analyzing kinetics of dLight signals with the ManM

We next analyzed the same raw data used to generate Figure 3 using the ManM. This method determines a %ΔF/F cutoff specific to each animal based on signal strength when animals are drug naïve, which is then maintained within animal for each drug condition. These cutoffs are determined during saline control sessions by matching each cutoff to the top of the band of noise that separates emergence of positive peaks from the baseline. This band of noise is evident in Figure 1 and specifically highlighted in Figures 2G **and 2H**. A simple t-test showed cocaine reduced the amplitude of signals compared to saline (t_2756_ = 7.071; saline: 0.0 ± 0.90%; cocaine: -9.0 ± 0.90 %; p<0.00001; Figure 4A) and increased signal width (t_2756_ = 7.878; saline: 0.0 ± 2.4%; cocaine: +34 ± 3.6%; p<0.00001; Figure 4B). These changes are supported by prior *ex vivo* studies with cocaine from our lab and others [27, 41, 62–65]. Because amplitude was decreased and width was increased, this altered the relationship between these variables. Slope was reduced by cocaine (t_6_ = 3.188; saline: 4.7 ± 1.4 ΔF/F(%)/s; cocaine: 2.5 ± 0.70 ΔF/F(%)/s; p=0.0189; Figure 4C). This shows that, at a given level of DA release, signals take longer to return to baseline (below the cutoff) after cocaine administration, indicating slower DA uptake.

**Figure 4.**
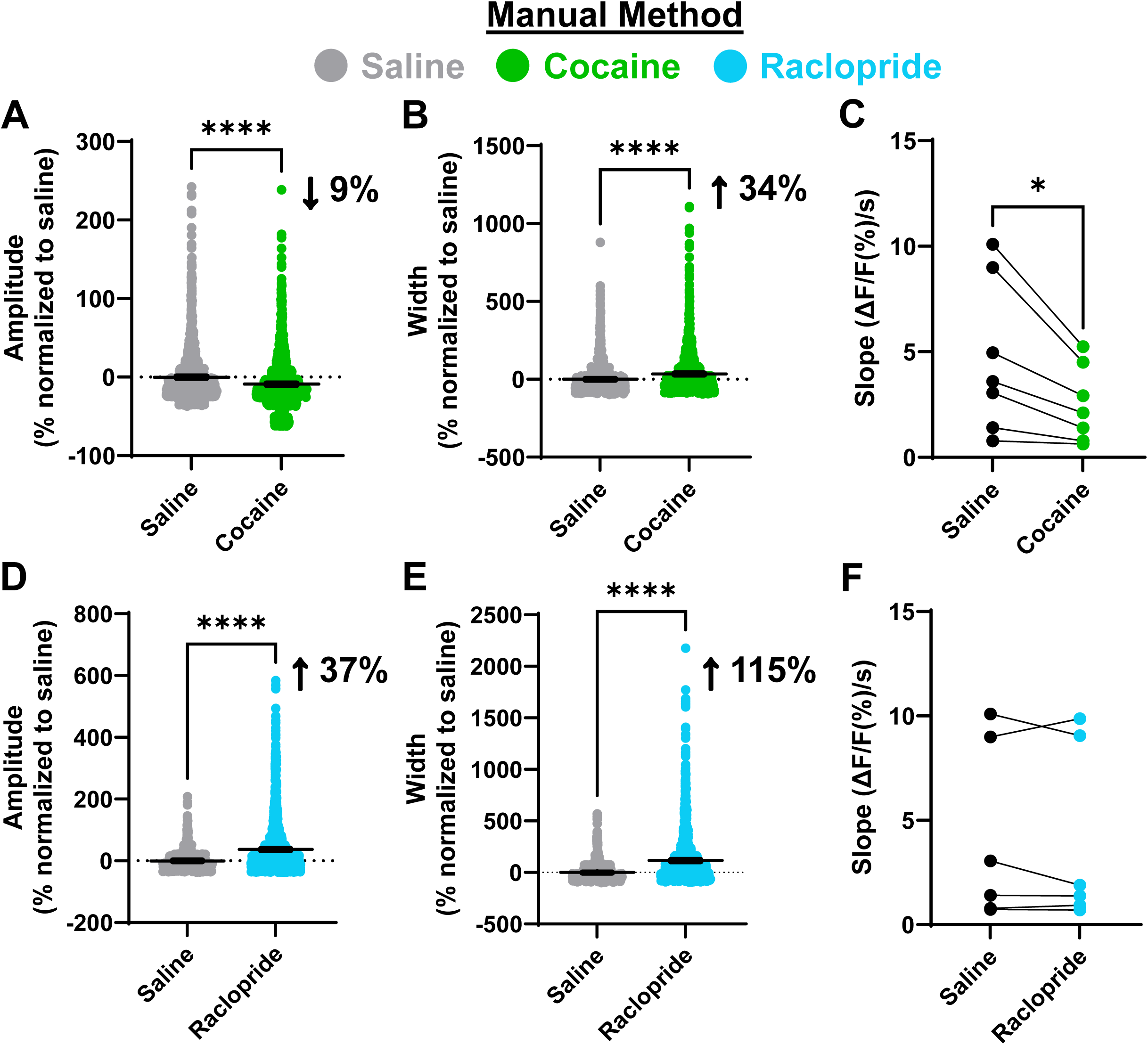
The Manual Method (ManM) shows that cocaine decreases DA uptake and raclopride increases DA release. The ΔF/F cutoff generated using the ManM was determined in each animal (range: 0.40 to 3.25%) and maintained between conditions. (A) Cocaine ( ● ) reduced signal amplitude and (B) increased signal width, (C) resulting in a reduction in the slope of their correlation, indicating slower uptake. (D-F) Using the ManM, raclopride ( ● ) increased both signal (D) amplitude and (E) width, resulting in no change in (F) slope, consistent with increased release. (*, p<0.05; ****, p<0.00001 (Bonferroni corrected); difference between saline and drug).

To validate the ManM, a separate, blinded investigator repeated the cocaine analyses. That analysis produced similar results, that cocaine reduced amplitude (t_4094_ = 3.493; saline: 0.0 ± 0.83%; cocaine: -4.3 ± 0.88%; p<0.001; not shown), increased width (t_4094_ = 9.620; saline: 0.0 ± 1.8%; cocaine: +33.2 ± 3.2%; p<0.00001; not shown), and reduced the slope of the correlation between amplitude and width (t_6_ = 2.814; saline: 4.4 ± 1.3 ΔF/F(%)/s; cocaine: 2.4 ± 0.66 ΔF/F(%)/s; p<0.05; not shown). Further, there was no difference in the primary hypothesized effect, an increase in signal width, between investigators (t_3038_ = 0.2256; p=0.8215; not shown). The results of the ManM also align with those of the FSCV dataset presented in **Figure S1**, that cocaine can reduce peak height of DA release, increase the time required for clearance, and alter the relationship between amplitude and width (slope). Overall, inputting %ΔF/F traces into pClamp, manually adjusting the baseline, and prescribing a %ΔF/F cutoff per animal (see **methods**) is a robust method to detect pharmacologically induced differences in the DA uptake rate of spontaneously arising DA signals using dLight.

In comparison to cocaine, raclopride markedly increased both signal amplitude (t_2917_ = 14.43; saline: 0.0 ± 0.90%; raclopride: +37.0 ± 1.8%; p<0.00001; Figure 4D) and width (t_2917_ = 15.52; saline: 0.0 ± 2.5%; raclopride: +115.2 ± 5.4%; p<0.00001; Figure 4E) using the ManM. Simple linear regression was run on all drug trials for all animals, followed by a paired t-test of slope values. Raclopride did not change the slope of the correlation compared to saline (t_5_ = 0.6568; saline: 4.2 ± 1.7 ΔF/F(%)/s; raclopride: 4.0 ± 1.7 ΔF/F(%)/s; p=0.54; Figure 4F). This suggests that, whereas cocaine inhibited DA uptake, raclopride disinhibited release, resulting in a proportional increase in signal amplitude and width. These findings also highlight the importance of assessing the relationship between these metrics, rather than solely reporting amplitude and width in isolation. Because it is difficult, if not impossible, to ascertain true DA concentrations *in vivo* when using fiber photometry, it is not feasible to determine an uptake rate based on concentration (like V_max_) using this technique. The validation of slope as an uptake metric is an invaluable tool for spontaneous dLight studies moving forward.

### 2.6. The ProM shows that smoothing and normalization can be used to detect signals in a standardized manner, if the preprocessed, non-normalized data are used to analyze amplitude and width

The issues with the analyses discussed thus far involve either blunting of the ability of the analysis to determine pharmacological effects using smoothing and normalization in the ZM or the need for different cutoff values to be used between animals (maintaining this cutoff within animals for drug comparisons) in the ManM. However, because a standardized cutoff can be used to *detect*, rather than analyze, signals using z-score normalization, we decided to use the Prominence Method (ProM) to detect signals and create a ‘timestamp’ for each signal, followed by assessment of the raw amplitude and width data that match those timestamps. We used a metric called “prominence” to determine which signals detected by z-scoring should be included.

Essentially, only dLight signals that were highly prominent, meaning larger than the baseline activity immediately surrounding the peak, were included.

Using the ProM, cocaine was shown to reduce signal amplitude (t_5350_ = 22.62; saline: 0.0 ± 0.97%; cocaine: -27.5 ± 0.74%; p<0.00001; Figure 5A) and increase signal width (t_5350_ = 5.314; saline: 0.0 ± 1.3%; cocaine: +12.1 ± 1.8%; p<0.00001; Figure 5B).

**Figure 5.**
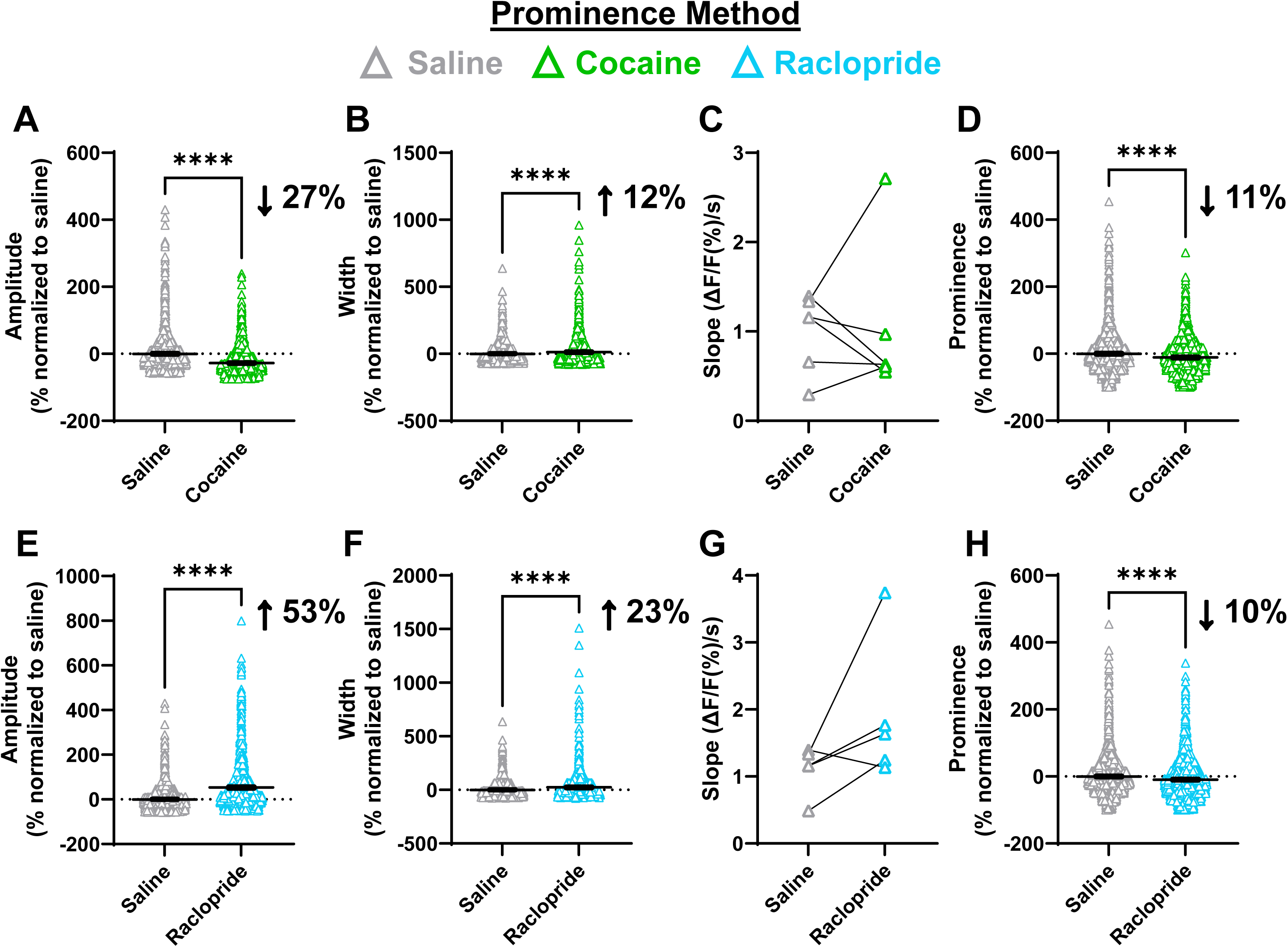
The Prominence Method (ProM) detects greater changes in signal amplitude but disrupts effects on slope. (A) Using the ProM with a cutoff of z = 1.0, cocaine ( Δ ) reduced signal amplitude, (B) increased signal width, (C) did not affect slope, (D) and decreased prominence. (E-H) Raclopride ( Δ ) increased both signal (E) amplitude and (F) width without affecting (G) slope, and reduced (H) prominence. (****, p<0.00001 (Bonferroni corrected); difference between saline and drug).

However, there was no effect on the slope (t_5_ = 0.05592; p=0.9576; Figure 5C). The ‘prominence’ metric was unique to the ProM, and cocaine reduced signal prominence (t_8570_ = 10.04; saline: 0.0 ± 0.82%; cocaine: -11.3 ± 0.75%; p<0.00001; Figure 5D), suggesting the basal DA activity below the cutoff was increased.

We next applied the ProM to assess effects of raclopride, which increased both signal amplitude (t_3858_ = 21.48; saline: 0.0 ± 0.998%; raclopride: +53.0 ± 2.52%; p<0.00001; Figure 5E) and signal width (t_3858_ = 7.827; saline: 0.0 ± 1.49%; raclopride: +23.3 ± 2.80%; p<0.00001; Figure 5F) without affecting slope (t_4_ = 1.815; p=0.1437; Figure 5G). Like cocaine, raclopride also reduced signal prominence (t_6640_ = 6.979; saline: 0.0 ± 0.868%; raclopride: -9.59 ± 1.06%; p<0.00001; Figure 5H). These results align with the ManM that raclopride increased DA release without affecting DA reuptake, as shown by proportional increases in amplitude and width resulting in no change in slope.

Overall, the ManM and ProM detected an increase in signal width after cocaine and an increase in DA release amplitude after raclopride. These primary hypothesized effects were absent using the ZM for analysis. Therefore, z-scoring can be used for signal detection as in the ProM, because the z-score phase is only used to tag or ‘timestamp’ signals, followed by retrieval of the non-normalized, preprocessed %ΔF/F data for analysis. A caveat is that the ProM was not able to detect a difference in frequency during the SCH23390 trials, suggesting that noise can still be amplified when there are no signals resultant from DA activation of the dLight receptor. However, unlike the ProM, the ZM lacks the ability to assess prominence for signal inclusion and to convert back to preprocessed values, making the ZM inadequate for any analysis of the kinetics of spontaneous signaling. As long as real spontaneous dLight signals are present, the ProM remains an invaluable tool for assessing changes in the peak height of DA release and signal width in a standardized, unbiased manner.

The relationship between DA release and time for clearance is significantly and positively correlated as reported herein using the ManM as well as in an *ex vivo* FSCV dataset, which was expected according to the literature [27–29]. However, use of the

ProM diminished the ability to detect differences in the slope of the correlation between amplitude and width during cocaine sessions. Using *in vitro* techniques, fluorescence intensity levels can be directly compared with a known DA concentration that is washed on during the experiment with consistent changes in %ΔF/F (for further information, see [5, 6, 8, 66]). Essentially, how ‘bright’ a signal can fluoresce in response to DA is known as its dynamic range [5, 13], but there are potentially confounding variables introduced in live animals. First, viral transduction efficiency depends on virus serotype [67–70], brain region [67, 70, 71], species [68, 69, 72], sex [73], age [74], route of delivery [70], and even variability between animals in the same species [75], including C57BL/6J mice [68]. Second, optic signals herein were sensed through a fiberoptic patch cable using the property of internal refraction [76, 77]. Light cannot escape *or* enter the core of the patch cable through the external jacket and cladding because they have a lower refractive index than the core [78], allowing light to propagate without degrading. This means that a fluorescent signal must be within the cone of acceptance and sufficiently close to the cannula to be propagated, factors that can differ between animals. We suggest that, while a given %ΔF/F value will not always equate to the same concentration of DA *between animals,* the same %ΔF/F value *within* an animal should consistently measure the same concentration of DA according to the sensor’s dynamic range.

While within-animal comparisons of changes in %ΔF/F work well according to the ManM, the same considerations do not apply when using z-scoring, as supported by the lack of effects shown with the ZM. Because signal prominence differs between pharmacological sessions, the z-score value that corresponds to a particular %ΔF/F value during one session or pharmacological condition does not necessarily equal the same %ΔF/F value from a recording taken on another day. Therefore, by taking amplitude and width values at potentially differing %ΔF/F values and DA concentrations, the slope of the correlation between amplitude and width is disrupted. In other words, because the ProM generates timestamps for analysis of amplitude and width during the z-scoring detection phase, this further suggests that slope is disrupted by normalizing entire traces. Thus, we suggest the ManM is superior to the ZM and ProM in detecting changes in width and slope, and is, therefore, best suited for detecting changes in DA uptake rates.

### 2.7. Significant differences occurred between methods in the ability to detect the primary hypothesized effects of cocaine and raclopride

To directly assess the ability of each method to detect changes in amplitude and width induced by cocaine and raclopride, we next performed a comparison between methods using the saline-normalized values for each drug. A series of one-way ANOVAs showed there was a significant effect of analysis method on relative change in amplitude (F_(2,4683)_ = 166.9; p<0.0001; Figure 6A) and width (F_(2,4683)_ = 39.58; p<0.0001; Figure 6B) during cocaine sessions. Post-hoc tests showed that the ProM detected the strongest decrease in amplitude compared to both the ManM and ZM (p<0.00001).

**Figure 6.**
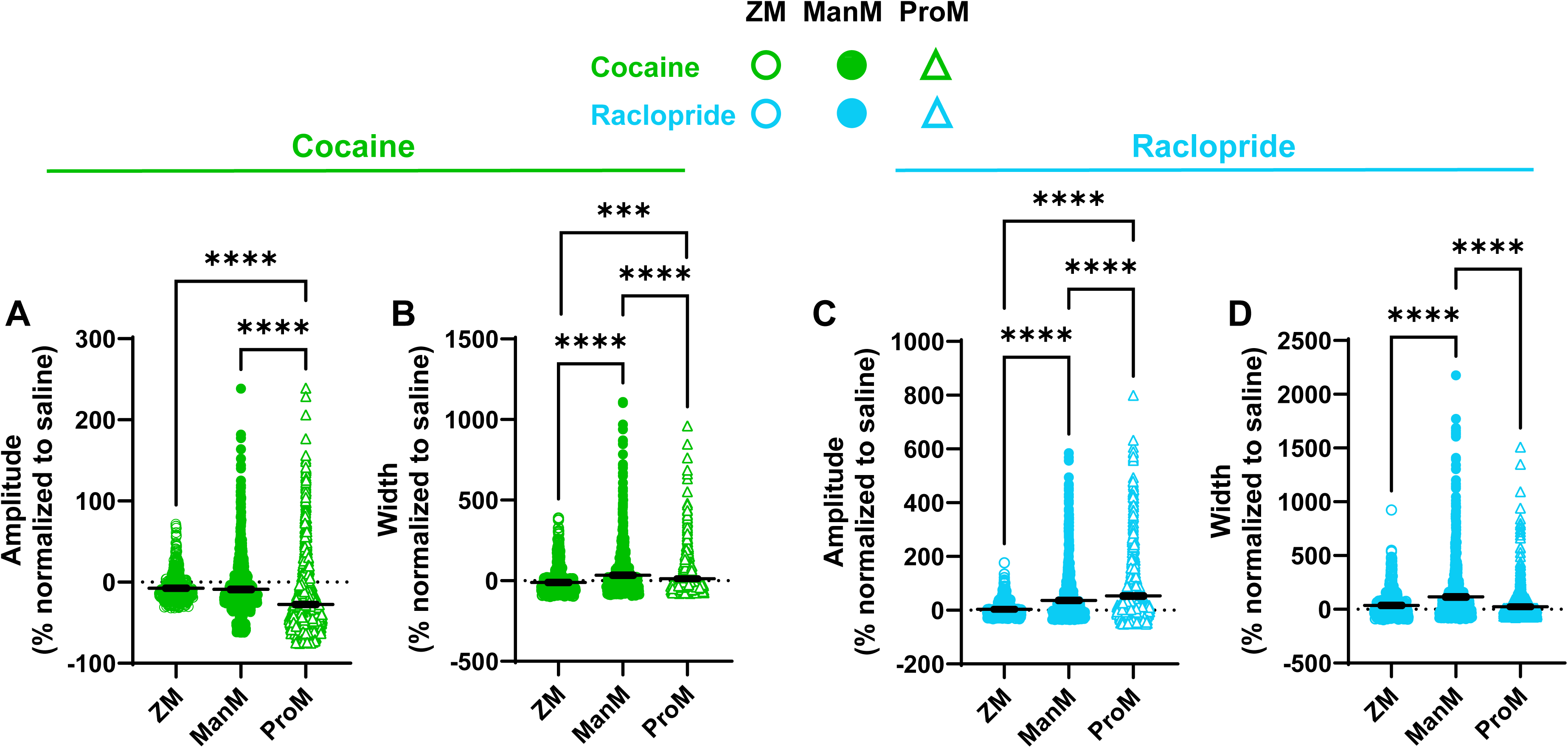
The Manual Method (ManM) and Prominence Method (ProM) were superior to the Z-score Method (ZM) in detecting effects of cocaine and raclopride. (A) Showing only the drug data normalized to saline control sessions, the ProM was superior to both the ManM and ZM ( ○ ) in detecting a reduction in signal amplitude induced by cocaine. (B) However, both the ManM ( ● ) and ProM ( Δ ) were superior to the ZM in detecting an increase in width, with the ManM performing the best. (C) Regarding raclopride, detection of an increase in amplitude was fully attenuated by the ZM ( ○ ), with the ProM ( Δ ) performing better than the ManM ( ● ). (D) In contrast, the ManM performed the best at detecting an increase in signal width. (***, p<0.0001 (Bonferroni corrected); ****, p<0.00001 (Bonferroni corrected); difference between methods).

Further, the ability to detect an increase in width induced by cocaine was fully attenuated using the ZM compared to the ManM (p<0.00001) and ProM (p<0.0001), whereas the ManM detected a larger cocaine-induced increase in width compared to the ProM (p<0.00001).

Regarding the ability to detect effects of raclopride, another series of one-way ANOVAs showed there was a significant effect of analysis method on relative change in amplitude (F_(2,4091)_ = 73.03; p<0.0001; Figure 6C) and width (F_(2,4091)_ = 127.2; p<0.0001; Figure 6D). Post-hoc testing showed significant attenuation of the ability to detect changes in amplitude using the ZM compared to the ManM and ProM (p<0.00001), whereas the ProM detected a larger increase by raclopride versus the ManM (p<0.00001). In contrast, the ManM detected the largest increase in width compared to the ZM and ProM (p<0.00001).

A summary of the considerations for each method are highlighted in **Table 1** and Figure 7. Overall, the ZM attenuated the ability to detect pharmacological effects of cocaine and raclopride. However, there was a different magnitude of effect between the

**Figure 7.**
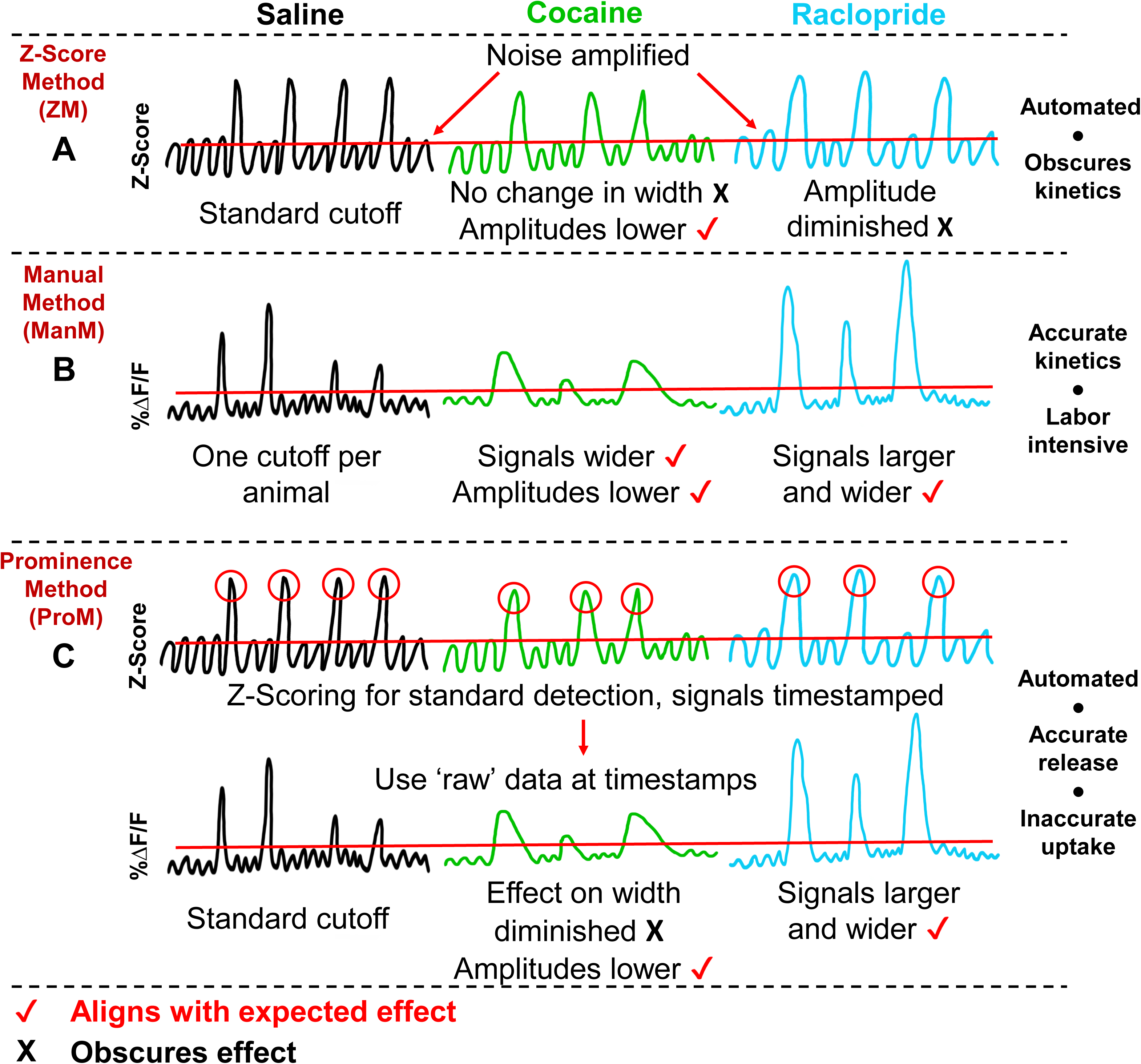
A summary of the outcomes of each method. A summary for each method is shown highlighting their ability, or inability, to accurately detect pharmacological effects of cocaine and raclopride. These include (A) the ZM, which obscures kinetic differences between drugs, (B) the ManM, which performs well for all drug conditions but is labor-intensive and time-consuming, and (C) the ProM, which is fast and automated with proper detection of changes in amplitude, but obscures effects on DA uptake.

**Table 1.** Highlights of each method.

|  | Pros | Cons | When to use | When not to use |
| --- | --- | --- | --- | --- |
| **ZM** | Fast & automated<br>•<br>Standardized cutoffs | May amplify noise<br>•<br>Masks effects on signal kinetics<br>•<br>Cannot be used on entire recordings taken longitudinally | With a narrow window of time set around behavioral events<br>•<br>When manipulations are applied *within a single recording* | To assess longitudinal changes<br>•<br>Anytime there is a change in processing over time – learning, experience, drug conditions, pathology, etc. |
| **ManM** | Accurate measurement of all kinetic measures<br>•<br>Superior assessment of <u>uptake</u> | Labor intensive<br>•<br>Different cutoff set for each animal<br>•<br>Requires blinding to experimental condition | Assess changes in spontaneous signal kinetics over time<br>•<br>Without *a priori* hypotheses on release and uptake |  |
| **ProM** | Fast & automated<br>•<br>Standardized cutoffs<br>•<br>Accurate measurement of amplitude, width<br>•<br>Superior assessment of <u>release</u> | Disrupts correlation between amplitude and width (uptake)<br>•<br>Can amplify noise during assessment of frequency | When release is the focus | When uptake is the focus |

ManM and ProM, depending on the variable. The ManM was better than the ProM at detecting changes in width for both drugs while, conversely, the ProM detected a larger change in amplitude than the ManM for both drugs. Above, we discussed that there was a reduction in prominence according to the ProM with both cocaine and raclopride, even though either drug had an opposite effect on amplitude. This means that the average DA activity below the defined cutoff point is larger with cocaine and raclopride, with the signals above the cutoff smaller in comparison to the ‘noise’. Because detected signal prominence is reduced by these drugs, this suggests that there are real differences in the background, or baseline, levels of DA that reflect real differences in tonic DA signaling rather than noise. This again precludes use of the ZM for analyzing spontaneous signaling.

### 2.8. A more stringent cutoff used for signal detection with the ProM increased the ability to identify effects of cocaine and raclopride to change signal amplitude and width

We wanted to ensure that the cutoff of z = 1.0 used with the ProM, which was more lenient than the cutoff used with the ZM, did not lead to a false conclusion that the ProM is always superior to the ZM. However, by rerunning the ProM using a more stringent cutoff of z = 2.6, the same cutoff used with the ZM, we showed that using preprocessed %ΔF/F values for analysis is superior to using z-scored values. Indeed, cocaine reduced signal amplitude (t_880_ = 14.59; saline: 0.0 ± 1.56%; cocaine: -29.9 ± 1.29%; p<0.00001; **Figure S2A**), increased width (t_880_ = 9.284; saline: 0.0 ± 2.59%; cocaine: +51.9 ± 5.11%; p<0.00001; **Figure S2B**), but did not change slope (t_4_ = 1.825; p=0.1421; **Figure S2C**). Further, raclopride increased both signal amplitude (t_783_ = 13.77; saline: 0.0 ± 1.66%; raclopride: +59.3 ± 3.79%; p<0.00001; **Figure S2D**), and width (t_783_ = 4.860; saline: 0.0 ± 2.93%; raclopride: +27.9 ± 4.74%; p<0.00001; **Figure S2E**) without affecting slope (t_4_ = 2.541; p=0.0639; **Figure S2F**). Overall, the ProM produced similar results using a cutoff of z = 1.0 (Figure 5) and z = 2.6 (**Figure S2)**, albeit with a different magnitude of effect. We did not analyze this for significance, but the average change of all amplitude and width values was larger with the more stringent cutoff. The most anticipated results, namely larger signal width due to cocaine and increased amplitude due to raclopride, were specifically increased the most. This supports that the cutoff used for the ZM was not the reason for attenuation of the ability to identify pharmacological effects, and that using z-scored values for analysis is inferior to using preprocessed %ΔF/F values as with the ManM and ProM.

Using the higher cutoff of z = 2.6 versus z = 1.0 with the ProM, 84% fewer signals were included in the saline – cocaine comparison, and 80% fewer were included in the saline – raclopride comparison. However, the results were the same with five times as many “smaller” signals included compared to using the z = 2.6 cutoff. This further suggests a difference in baseline variability between drug conditions that can affect the results, given that results were similar using a much lower z-score cutoff but using preprocessed %ΔF/F values for analysis. Therefore, rather than ‘noise’, it appears likely that signals around a lenient z-score cutoff of z = 1.0 are resultant from DA activation of the dLight receptor. Overall, the ProM is the most useful method in detecting differences in signal amplitude, particularly when a reduction in signal amplitude is hypothesized. Another major benefit of this method is that a standard z-score and prominence cutoff can be applied to all animals and sessions. In contrast, with the ManM, a cutoff must be prescribed for each animal, though this is still maintained between pharmacological conditions, and any dLight signals that are sufficiently reduced by a given manipulation would be excluded with the definitive cutoff.

## 3. Conclusions

This paper assessed the same fiber photometry dataset using three distinct methods to determine the best way to analyze spontaneously arising DA release events. The basis of each method was to apply a cutoff, above which the amplitude, width, and frequency of signals were assessed. We determined that the use of smoothing to remove baseline drift and z-scoring to normalize traces and apply a standard cutoff with the Z-Score Method was not appropriate for the final analysis. This is because data were taken longitudinally, and z-scoring attenuated variability of experimental interest when manipulations were given between recordings. This decreased the ability to identify the effects of D1R antagonism to block signal frequency, of DAT inhibition to increase signal width, and of D2R antagonism to increase DA release amplitude. In contrast, the Manual Method analyzed non-normalized, preprocessed data with a %ΔF/F cutoff, which performed better at detecting pharmacologically induced changes in amplitude, width, and frequency. The major limitations included that baseline drift was adjusted manually, drastically increasing the time required for analysis and leaving room for experimenter bias, and different cutoffs were applied between animals based on dLight expression. Therefore, we developed a final method (the Prominence Method) that used smoothing and z-score-based normalization solely for signal identification, followed by use of the original, non-normalized data for analysis and an assessment of prominence to determine if signals were sufficiently separated from the baseline to be included. This allowed for standard z-score and prominence cutoffs to be used between all conditions when baseline variability differed. The major drawback was that a given z-score likely does not reflect the same %ΔF/F value between conditions, disrupting the assessment of DA uptake using the slope of the correlation between amplitude and width. In conclusion, z-scoring is not appropriate for the quantification of spontaneous signals. The Manual Method was the best at detecting changes in DA uptake rate, and the Prominence Method was superior at detecting changes in DA release. Understanding these different types of analyses for fiber photometry data and the circumstances under which each is best used will advance the field towards a common set of analyses that will standardize results and make comparisons across data sets and labs much easier and more accurate.

## 4. Methods

### 4.1. Animals

Animal protocols described herein were reviewed and approved by the Wake Forest University Institutional Animal Care and Use Committee. Seven-week-old C57BL/6J mice (n = 12 male, n = 10 female; Jackson Laboratory, Bar Harbor, ME) were allowed to acclimate in our colony room for one week housed on a 12:12 light/dark cycle (lights on: 0600, lights off: 1800). They received water and standard rodent chow *ad libitum*. Mice were approximately 12 weeks old at the start of testing.

### 4.2. Stereotaxic surgery

Surgeries were previously described in detail [46]. Briefly, mice (n = 5 male, n = 5 female) were anesthetized and received 700 nL injections of the dLight virus (AAV5- hSyn-dLight 1.2: Addgene, Catalogue No.: 111068-AAV5) into the nucleus accumbens (NAc) core (AP: +1.1 mm, ML: ± 1.3 mm, DV: +4.6 mm; from Bregma), counterbalanced unilaterally between animals. A fiber optic cannula was then lowered to the injection site (DV: +4.5 mm from Bregma). The surgical site was closed and cannula locked into place with adhesive luting cement (Parkell C&B-Metabond, Part No.: S380) and black dental cement (Lang Dental, Part No.: 1530 & 1404). Pain management was delivered preoperatively and for ≥ 2 days postoperatively until recovery.

### 4.3. dLight fiber photometry acquisition

After approximately four weeks of postoperative recovery and viral expression of dLight, mice were tested in their home cages to establish excitation intensity of the 465 nm light required to observe dLight signals. Photometry recordings were taken in the same manner as described previously [46]. Briefly, the Fiber Photometry Systems (RZ10X processor, Synapse software) setup from Tucker Davis Technologies (Alachua, FL), with supporting wiring from Doric Lenses (Quebec, Canada), was used to modulate two excitation wavelengths, the 405 nm (isosbestic control) and 465 nm (dLight- dependent) lights. LEDs were channeled into 0.57NA, 400μm core pre-bleached, low-autofluorescence patch cords with intensities maintained throughout the study at the tip of the patch cord at 15 μW for the 405 nm light and 15-30 μW for the 465 nm light (maintained within animal). Synapse was used to measure, in real time, demodulated and low-pass filtered (6 Hz) transduced fluorescence signals via the RZ10X.

### 4.4. Pharmacology and home cage testing

Mice were previously acclimated to dLight recordings and injections during initial home cage testing to determine intensity levels for 465 nm excitation. On experimental days, they were brought to the behavioral space and allowed to acclimate for at least one hour. They were administered either saline (IP, time matched to drug trials), cocaine (IP, 10 mg/kg, immediately before dLight recording), raclopride (IP, 0.3 mg/kg, 15 minutes prior), or SCH23390 (IP, 1 mg/kg, 30 minutes prior). Mice were then tethered to patch cords and the dLight recording was started. They were allowed to explore the home cage, with enrichment and cage tops removed, for 10 minutes. Drugs were administered according to a Latin square design, except SCH23390, which was given last to prevent attenuation of dLight signals.

Cocaine ((-)-cocaine HCl, RTI Log No.: 14201-134) was generously provided by the NIH and mixed in saline at 2 mg/mL. Raclopride (S(-)-raclopride (+)-tartrate salt, CAS: 98185-20-7) was purchased from Sigma Aldrich (Product No.: R121) and mixed in saline at 0.12 mg/mL. SCH23390-HCl (CAS: 125941-87-9) was purchased from Tocris (Product No.: 0925) and mixed in saline at 1 mg/mL.

### 4.5. dLight fiber photometry analysis

Raw Synapse files were extracted into a format readable by MATLAB (v. R2023a). Raw fluorescence values resulting from light excitation were preprocessed to remove bleaching and motion artifacts by subtracting the isosbestic control from the experimental channel. This resulted in a %ΔF/F trace for entire recording sessions.

For the Z-Score Method, we used code developed by Martianova et al. [79] and Zhang et al. [80] in MATLAB. We refer you to these studies for a detailed discussion of the statistics underlying this code, which was designed to remove drift inherent in analytical datasets and amplify potential peaks. Here, we applied this code to a much different type of dataset. Briefly, this method first applies a smoothing filter to the isosbestic 405 nm-excited and experimental 465 nm-excited recordings, fits them using linear regression, normalizes the traces into z-score values, and then subtracts the isosbestic from the experimental channel. This is accomplished using the adaptive iteratively reweighted Penalized Least Squares (airPLS) algorithm. This algorithm compares every %ΔF/F value to the values that occurred immediately before and after and normalizes the whole signal to z-score values based on the variability inherent within each recording. This serves at least two functions, including correcting the baseline drift as well as generating a list of z-scores by which to analyze signals. We set a z-score threshold of z = 2.6 (p<0.01), which is a relatively strict cutoff intended to exclude any noise from being analyzed. We applied this cutoff to all recordings from all animals and analyzed signals to determine amplitude (the peak z-score), width (the time it took for the signal to rise above then back below the cutoff), and frequency (peak-to-peak in signals/s).

In a separate set of analyses (Manual Method – ManM), we adjusted for bleaching and motion artifacts by subtracting the isosbestic control from the experimental signal. Next, %ΔF/F traces, which were not normalized further, were loaded into AXON pCLAMP software (Clampfit v. 11), and baselines were manually adjusted for drift. A vehicle control trial from each animal was first assessed to determine the %ΔF/F cutoff value, which was specific for each animal and maintained for cocaine, raclopride, and SCH23390 trials. Cutoffs were determined visually during drug-naïve, saline control trials by matching each cutoff to the top of the band of noise that separates where individual positive peaks emerge from the background. This band of noise is visually evident in Figure 1 and specifically highlighted in Figures 2G **and 2H**. Cutoffs were also matched visually above this band of noise to the amplitude of the negative peaks, which typically have a much smaller change from baseline. Cutoffs for all drug trials were maintained within animals from the cutoff determined during vehicle control trials that occurred within one week of the drug trial. pClamp was used to generate the amplitude (the maximal %ΔF/F value, including the distance from a baseline of %ΔF/F = 0 to the peak), width (the time it took for the signal to exceed then fall back below the cutoff), and frequency (peak-to-peak in signals/s) of signals.

In the final method (Prominence Method - ProM), we adjusted for bleaching, drift, and motion artifacts through normalization by detrending the 465-nm-excited signal and subtracting a moving average smoothed (width=100 s) version of the signal. This signal was then z-scored and a z-score threshold was applied to systematically *detect* signals. We decided to use a more permissive cutoff of z = 1.0 (p=0.1587) in the main results, because a second metric – prominence – was used to determine signal inclusion (see below). Signals detected in this manner were used to create ‘timestamps’ for each peak, which were used to find the corresponding signals in the non-normalized %ΔF/F traces. The non-normalized, preprocessed %ΔF/F signals at the same timestamps were then quantified for amplitude (the maximal %ΔF/F value), width (the time it took for the signal to exceed then fall back below the cutoff), and frequency (peak-to-peak in signals/s) of signals. After this, each timestamped signal was assessed for its prominence. This considers that, on different recording days with either vehicle or drug injections, the variability in signals *below* the cutoff may be affected (which could affect the z-score calculation and signal detection). Accordingly, each signal was assessed for prominence, the difference between the signal’s peak height and the level of activity that occurs immediately around it. This allows a fairer inclusion criteria than applying a blanket cutoff between recording days, which occurs in the ZM and ManM. Signals that were included after this second ‘prominence cutoff’ were analyzed and presented. We used a prominence value of 2 for inclusion criteria, according to the findpeaks.m function within MATLAB. Finally, we plotted each signal 1 s before and 1 s after the identified peak using the timestamps generated in earlier steps and present the average traces of spontaneous signals in vehicle versus drug trials.

### 4.6. Fast-scan cyclic voltammetry (FSCV)

The general methods used within our lab for FSCV have been described extensively [35–40]. Briefly, mice (n = 7 male, n = 5 female) were anesthetized and decapitated, and brains were transferred to ice cold, oxygenated artificial cerebrospinal fluid (aCSF). Brains were sliced into 300 µm coronal slices and allowed to equilibrate in the recording dishes for ≥ 1 hour. Within recording dishes, aCSF was maintained at 32⁰C and a flow rate of 1 mL/min. A stimulating electrode was placed in the NAc core next to a recording electrode, cycling at 10 Hz, and a single monophasic electrical pulse (750 µA, 4 ms) was applied every 300 seconds until signal stabilization (<10% change in peak height) for at least three measurements. After stabilization, cocaine was applied to the aCSF in cumulative doses (0.3, 1, 3, 10, 30 µM), with seven stimulations applied at each dose. Only data at the highest dose of cocaine (30 µM) were shown as a proof of concept for dLight analyses. FSCV data were recorded and analyzed using Demon Voltammetry [27].

### 4.7. Statistical analysis

The amplitude (%ΔF/F or z-score) and width (ms) of signals from pharmacological challenges were normalized to vehicle control trials within each animal. These values were then compared using simple t-tests. Frequency (signals/s) was assessed using paired t-tests, and the effect of analysis method on frequency was determined using one-way analysis of variance (ANOVA) with Tukey’s multiple comparisons post-hoc test to determine group differences. To generate the ‘slope’ variable, simple linear regression was conducted on amplitude versus width values for all signals within each recording session, followed by a paired t-test on the resultant slope values. One-way ANOVA was used to compare the relative change in amplitude and width values during drug conditions between the three analysis methods, with Tukey’s multiple comparisons test used for post-hoc testing. For the FSCV data in **Figure S1**, simple linear regression and Pearson correlation were used to show a relationship between peak height of DA release and the T_80_ metric used to show the time required for clearance. Paired t-tests were used for the before-and-after comparison of release and clearance data during baseline versus cocaine application within the same brain slice. Statistical analysis was conducted via GraphPad Prism (v. 10.1.2). The direction of change for amplitude and width variables was the same between sexes for cocaine and raclopride trials, so sexes were combined for analyses.

To provide further validity of the statistical results, different alpha values were used to determine significance based on the comparisons being performed. An average value for each animal was required to fairly compare the slope and frequency data between drugs and methods. This was because slope was generated from the correlation between amplitude and width, and only one value was generated per session. Further, using the ManM, there were no detected signals in some trials during the SCH23390 condition, meaning that using all values like in the amplitude and width comparisons would skew results when some sessions had no detected values. Thus, there was a relatively small sample size (n = 5 to n = 7 animals depending on the drug or method tested). There was also a relatively small sample size for the FSCV data (n = 13 animals, n = 29 slices). Therefore, a standard alpha value was used, yielding p = 0.05 as a threshold for significance, with a standard pattern of symbols to indicate significance within figures (*, p<0.05; **, p<0.01; ***, p<0.001; ****, p<0.0001). However, there were far more amplitude and width values included for comparison (n = 100s to n = 1,000s of signals depending on the drug or method tested). Therefore, we conducted a Bonferroni correction, dividing the standard α = 0.05 by the number of overall comparisons, or subfigures, within each main figure. This yielded a significance threshold of p = 0.0125. For clarity in presenting the results, we rounded down, resulting in the following pattern to indicate significance (*, p<0.01; **, p<0.001; ***, p<0.0001; ****, p<0.00001).

## Safety

There are no safety hazards or risks associated with this report.

## Abbreviations

DA, dopamine; FP, fiber photometry; ZM, Z-Score Method; ManM, Manual Method; ProM, Prominence Method; D1R, dopamine D1-like receptor; DAT, dopamine transporter; D2R, dopamine D2-like receptor; FSCV, fast-scan cyclic voltammetry

## Corresponding Author Information

Sara R Jones

Department of Translational Neuroscience Wake Forest University School of Medicine Medical Center Blvd

Winston-Salem, NC, 27157

## Author Contributions

Conceptualization: SRJ, KMH, CWW; Methodology: CCL, CWW, SWC, SRJ, KMH; Software: CWW, CCL, SWC; Validation: CWW, CCL, SWC, SRJ, KMH, CYS, RS;

Formal Analysis: CWW, RS; Investigation: CWW, CYS; Resources: SRJ, CCL, KMH,

SWC; Data Curation: CWW; Writing – Original Draft: CWW; Writing – Review and Editing: SRJ, KMH, CCL, CWW, SWC; Visualization: CWW, SRJ, KMH, CCL; Supervision: SRJ; Project administration: SRJ, KMH, CWW; Funding acquisition: SRJ, KMH.

## Acknowledgement

We would like to thank the Integrative Neuroscience Initiative on Alcoholism and Dr. Alex Deal for their insights during the development of our methods and story. We would also like to thank the National Institute on Alcohol Abuse and Alcoholism as well as the National Institute on Drug Abuse for the funding that enabled this work. Finally, we thank the Wake Forest University School of Medicine for hosting the materials, equipment, animals, and other resources required for the completion of this work.

---

**Figure S1.**
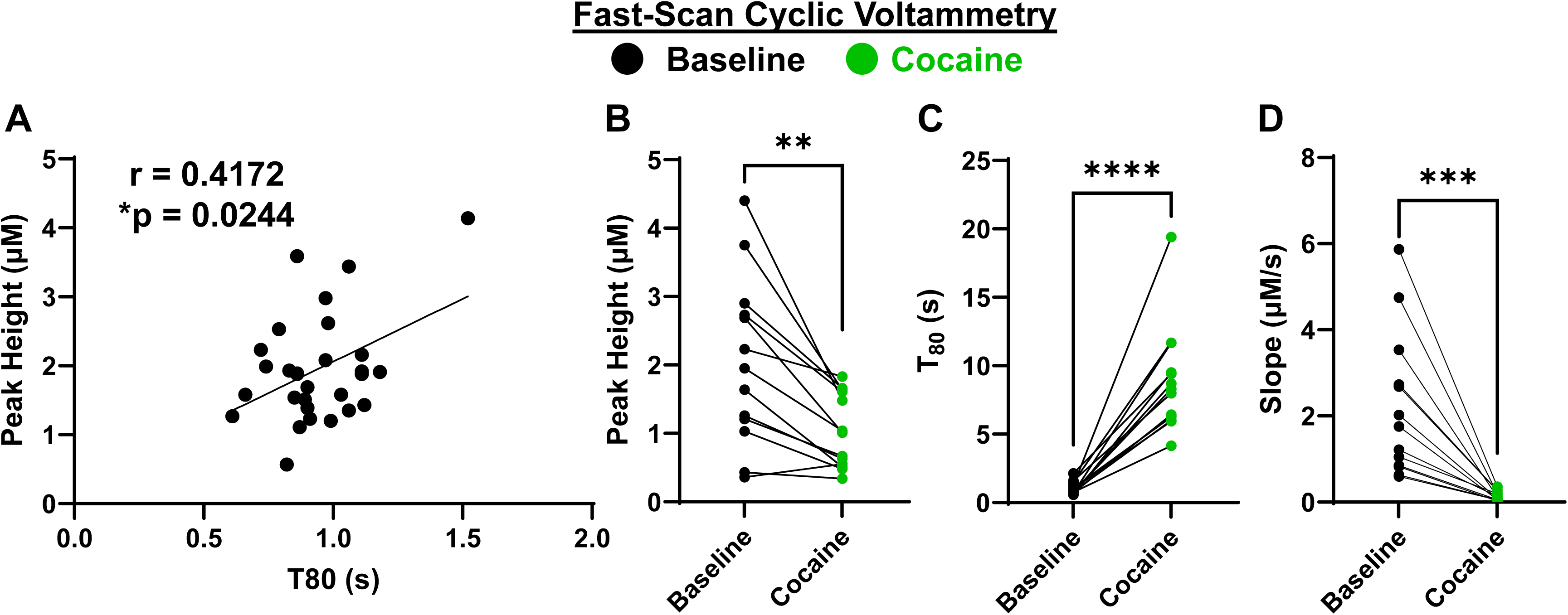
Fast scan cyclic voltammetry (FSCV) confirms the principles of spontaneous dLight analysis. A primary hypothesis underlying the dLight analyses herein of spontaneous signaling is that the peak height of DA release is intimately tied to the amount of time needed for signal clearance. **(A)** Indeed, using FSCV, peak height of stimulated DA release in NAc core-containing brain slices was correlated with the amount of time it took for clearance of 80% of the signal (T_80_) at baseline ( ● ). **(B-C)** At a high dose of cocaine (30 µM) ( ● ), **(B)** the peak height of DA release was reduced and **(C)** T_80_ was increased. **(D)** Finally, we performed the same “slope” comparison similarly to the FP data. Slope (peak height/T_80_) was reduced by cocaine. Therefore, effects of cocaine shown using FSCV align with the effects of cocaine herein when using the ManM. (*, p<0.05; significant correlation) (**, p<0.01; ***, p<0.001; ****, p<0.0001; difference between baseline and cocaine).

**Figure S2.**
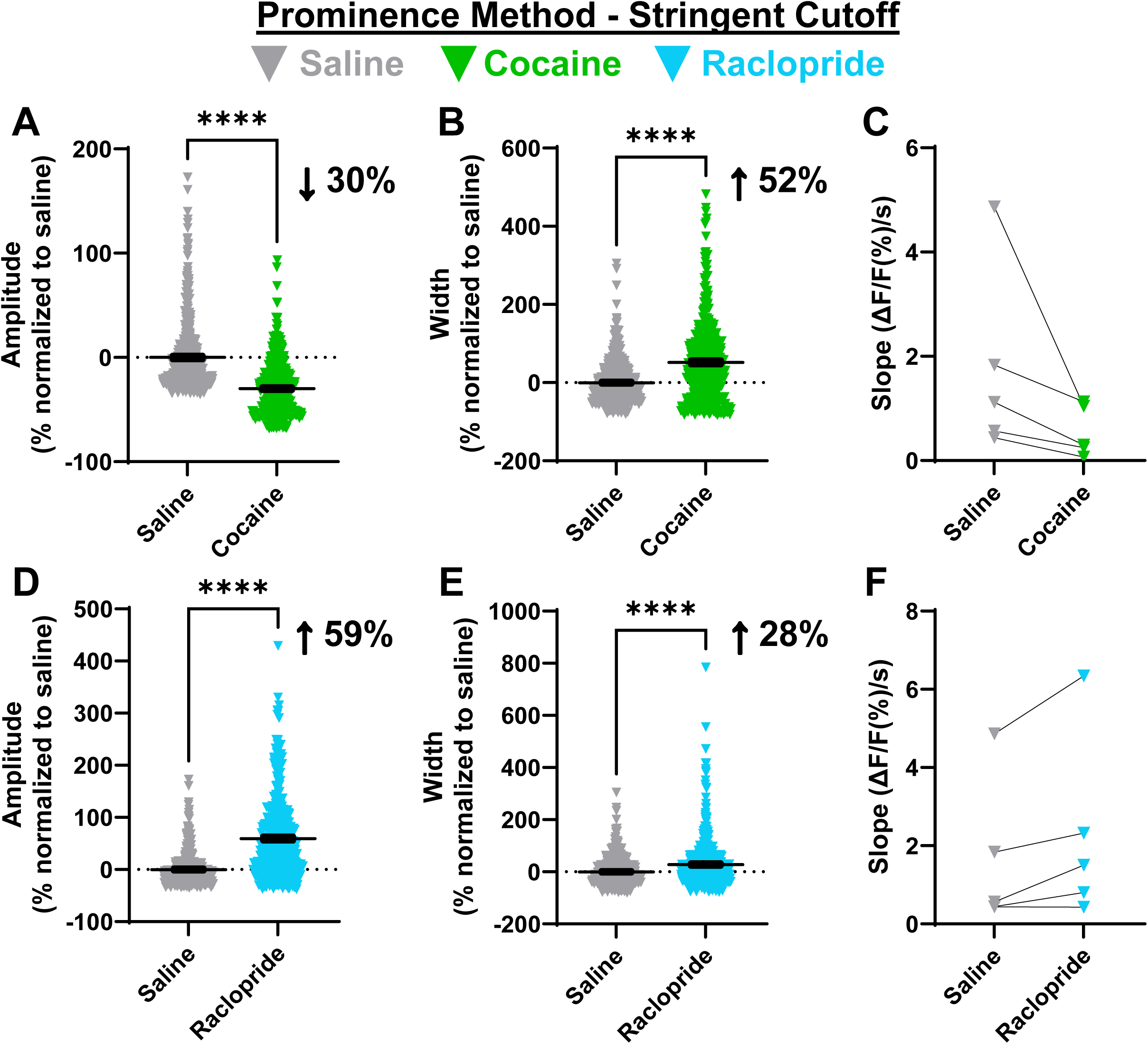
The Prominence Method (ProM), with a more stringent cutoff, promotes detection of primary hypothesized effects. The ProM was used with a more stringent cutoff for detection (z = 2.6), which was the same cutoff used with the ZM, to show that z-scoring was the issue and not the specific standardized cutoff. Similar to results using z = 1 (Figure 5), **(A-C)** cocaine ( ▼ ) **(A)** reduced signal amplitude and **(B)** increased signal width, **(C)** without affecting slope**. (D-F)** Raclopride ( ▾ ) was shown to **(D)** increase signal amplitude and **(E)** width, **(F)** without affecting slope. (****, p<0.00001 (Bonferroni corrected); difference between saline and drug).

